# The role of meaning in visual memory: Face-selective brain activity predicts memory for ambiguous face stimuli

**DOI:** 10.1101/366641

**Authors:** Timothy F. Brady, George A. Alvarez, Viola S. Störmer

**Author notes:** Please address correspondence to: Timothy F. Brady, Department of Psychology, University of California, San Diego, McGill Hall, Room 5322, 9500 Gilman Dr. #0109, San Diego, CA 92093.

## Abstract

How people process images is known to affect memory for those images, but these effects have typically been studied using explicit task instructions to vary encoding. Here, we investigate the effects of intrinsic variation in processing on subsequent memory, testing whether recognizing an ambiguous stimulus as meaningful (as a face vs. as shape blobs) predicts subsequent visual memory even when matching the perceptual features and the encoding strategy between subsequently remembered and subsequently forgotten items. We show that single trial EEG activity can predict whether participants will subsequently remember an ambiguous Mooney face image (e.g., an image that will sometimes be seen as a face and sometimes not be seen as a face). In addition, we show that a classifier trained only to discriminate between whether participants perceive a face vs. non-face can generalize to predict whether an ambiguous image is subsequently remembered. Furthermore, when we examine the N170, an ERP index of face processing, we find that images that elicit larger N170s are more likely to be remembered than those that elicit smaller N170s, even when the exact same image elicited larger or smaller N170s across participants. Thus, images processed as meaningful – in this case as a face–during encoding are better remembered than identical images that are not processed as a face. This provides strong evidence that understanding the meaning of a stimulus during encoding plays a critical role in visual memory.

**Significance Statement:** Is visual memory inherently visual or does meaning and other conceptual information necessarily play a role even in memory for detailed visual information? Here we show that it’s easier to remember an image when it’s processed in a meaningful way -- as indexed by the amount of category-specific brain activity it elicits. In particular, we use single-trial EEG activity to predict whether an image will be subsequently remembered, and show that the main driver of this prediction ability is whether or not an image is seen as meaningful or non-meaningful. This shows that the extent to which an image is processed as meaningful can be used to predict subsequent memory even when controlling for perceptual factors and encoding strategies that typically differ across images.

## Introduction

If you are asked to remember the visual details of an image – say, which exact face you saw – do you encode this information in a purely perceptual format, like a camera would store a photograph? Or does human memory always – even for visual details – depend on how meaningful that stimulus is to a given person? Memory for visual details is often quite impressive (Brady et al. 2008; Brady et al. 2013; Hollingworth, 2004), suggesting that detailed visual information is frequently stored in visual long-term memory. At the same time, it has long been argued that memory is semantically organized (Collins & Loftus, 1975), and that both elaborative encoding and strong retrieval cues are critical to the success of long-term memory. A meaningful interpretation of an image may allow for a more elaborative initial encoding, creating more or stronger paths to access the memory (Bradshaw & Anderson, 1982; Bower, Carlin & Dueck, 1975).

Consistent with the idea that how a stimulus is initially processed is critical for memory performance, a large body of research has shown that neural activity patterns during encoding can predict subsequent memory (Wagner et al., 1998; Kuhl, Rissman, & Wagner, 2012; Sanquist et al., 1980; Paller & Wagner, 2002; Daselaar, Prince, & Cabeza, 2004). However, for the most part this existing work does not distinguish whether such encoding-related activity is a result of differences in perceptual input, differences in encoding strategies, or differences in how information connects to meaningful concepts or semantic knowledge. In particular, given that in the majority of memory tasks the same items tend to be remembered by each person (Isola et al. 2011, Bainbridge, Isola, & Oliva, 2013), in an experiment examining the neural correlates of subsequent memory performance, the stimuli that are subsequently remembered are likely to differ along a number of dimensions from those that are subsequently forgotten. Some work has specifically focused on the neural correlates of meaningful processing in subsequent memory. For example, when participants are instructed to make semantic (vs. not) judgments of a stimulus at encoding, neural processing at encoding differs between these two conditions, and these differences predict subsequent memory performance (Sanquist et al., 1980; Dubé et al., 2013; Hanslmayr, Spitzer, Bäuml, 2008; Hanslmayr & Staudigl; 2014). However, despite examining the role of meaning and meaningful processing, this work involves an explicit manipulation of encoding strategy to require participants to focus on this aspect of the stimuli. Thus, the neural results likely reflect the performance of a different task by participants, rather than the particular processing of that stimulus that results in the most robust memory formation.

In the present study we test whether subsequent visual memory can be predicted by the extent to which a stimulus is initially processed in a meaningful way, independent of perceptual features, familiarity, or differences in encoding strategies. We use a set of visual stimuli –ambiguous Mooney images (Mooney, 1957) – that are carefully matched such that different images tend to be remembered by different people, with no reliable predictability in which images are remembered. To examine whether subsequent memory performance can be predicted based on how the stimulus is processed initially, we record the brain’s electrophysiological activity during encoding of these stimuli and test whether single-trial EEG activity at encoding can predict whether a stimulus is later remembered or forgotten. In addition, we make use of a neural signature that is selective to face processing: the N170 component (George, Jemel, Fiori, Chaby, & Renault, 2005; Hadjikhani, Kveraga, Naik, & Ahlfors, 2009; Liu et al., 2014), which allows us to assess whether a stimulus was likely seen as a face or not at encoding. Overall, our data suggests that memory performance is significantly affected by how meaningful – in this case face-like– the stimulus was processed at encoding, even when the stimulus input and the encoding task is held constant across stimuli and individuals.

## Materials and Methods

### Participants

Data from 24 participants were included in the final data analysis; data from three additional participants had to be excluded because less than 15 trials were available in at least one of the conditions due to artifacts in the EEG. All participants had normal or corrected-to-normal vision and were between 18 and 28 years of age. All studies were approved by the Institutional Review Board and all participants gave written informed consent prior to the experiment.

### Experimental Design

Participants were asked to remember a sequence of two-tone images ("Mooney images"; Mooney, 1957; see Figure 1). Stimuli consisted of 140 clear, unambiguous Mooney faces (stimuli designed so all participants would see them as a face), 140 ambiguous Mooney faces (stimuli designed so some participants would see them as a face and some would not see them as a face), and 140 non-face images. The unambiguous and ambiguous face images were chosen by using rankings collected on independent naive individuals indicating whether they saw each stimulus as a face or not as a face (Schwiedrzik, Melloni, Schurger, 2018). The non-face images were created by segmenting continuous regions of black and white from the face images and scrambling these parts by placing them at random locations and random angles within a new image, thus preserving overall luminance and the shape of each of the regions while removing the holistic connection between regions that allows the images to be seen as faces. Half of the images of each set were randomly selected for each participant to be presented during the study blocks, and the remaining half of the images served as foils during the test block. During the study blocks, participants were shown a stream of images on a gray background. Each image was presented for 500 ms in the center of the screen, followed by a 1000ms fixation cross. Participants were instructed to remember the images while keeping their eyes in the center of the screen. Participants were shown 210 images overall, divided into 10 study blocks (21 images per block, seven of each stimulus type). After each study block, participants performed a memory test by making an old/new judgment: Forty-two images were presented one at a time, and participants had to indicate for each test image whether they had seen it in the previous block ("old") or not ("new") by pressing one of two buttons on a keyboard. Half of the test images were old (i.e., they were presented during the previous encoding block), and the remaining half of the images were new (i.e., foils; participants had not seen them before). Study images from each stimulus set were selected at random for each participant and presented in random order. Participants were instructed to emphasize accuracy, not speed, in making their judgments.

**Figure 1.**
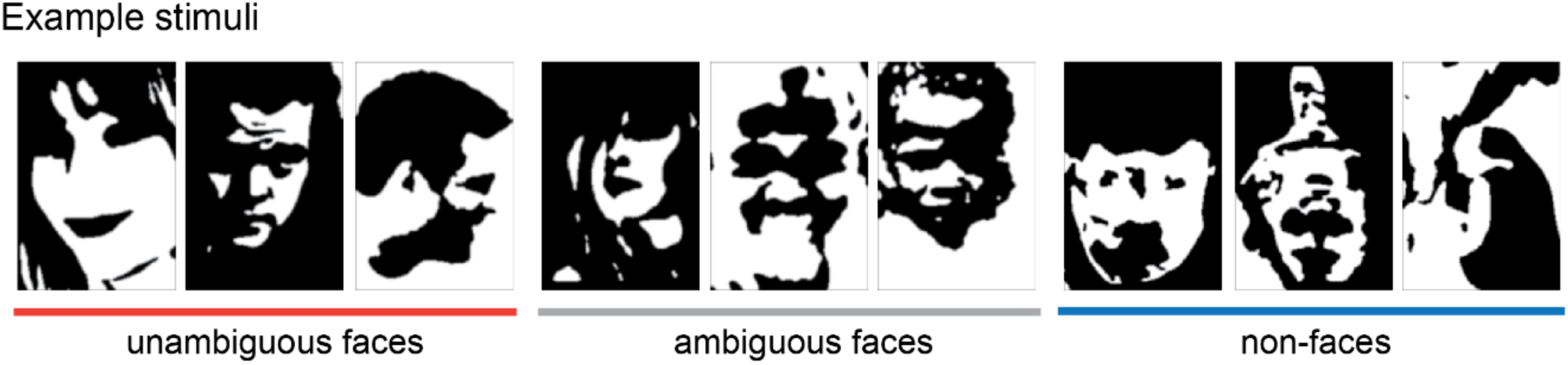
Left: Unambiguous Mooney face stimuli; Center: Ambiguous stimuli that some participants see as faces whereas others do not see as faces (when briefly presented); Right: Non-faces designed to match the face stimuli in low-level characteristics; made from scrambling and inverting parts of the unambiguous and ambiguous faces.

### Electrophysiological Recordings

While participants performed the memory task, brain activity was recorded continuously from 32 Ag/AgCI electrodes arranged according to the 10–20 system, mounted in an elastic cap and amplified by an ActiCHamp amplifier (BrainVision LLC). The horizontal electro-oculogram was acquired using a bipolar pair of electrodes positioned at the external ocular canthi, and the vertical electro-oculogram was measured at electrode FP1, located above the left eye. All scalp electrodes were referenced to an electrode on the right mastoid online, and were digitized at a rate of 500 Hz. Signal processing was performed with MATLAB (The MathWorks) using the EEGLAB and ERPLAB toolboxes (Delorme & Makeig, 2004; Lopez-Calderon & Luck, 2014) and custom-written scripts. Continuous EEG data was filtered offline with a bandpass of 0.01-112 Hz. Trials with horizontal eye movements, blinks, or excessive muscle movements were excluded from the analysis. Artifact-free data was re-referenced to the average of the left and right mastoids. For all further analysis, including decoding, data were epoched into trials and baselined to the 200ms pre-stimulus interval, and only data from trials without artifacts were included in the analysis.

### Statistical Analysis

#### Behavioral Analysis

To assess memory independent of response bias we calculated d’ as our measure of sensitivity. We then used a repeated-measures analysis with image type as a within-subject factor, followed by planned t-tests to assess the extent to which memory differs between our three conditions.

#### Decoding Analysis

Our main electrophysiology analysis is based on a Support Vector Machine (SVM) decoding approach (e.g., Cox & Savoy, 2003). In particular, we use EEG activity from each encoding trial to train a classifier to distinguish between the conditions and to predict subsequent memory. The input feature vector in each case consists 806 features: 31 electrodes x 26 time points (a running average of 20ms, sampled every 20ms between 100ms and 600ms post-stimulus). For situations where we train and test on the same conditions, we then apply a leave-one-out classification procedure with 500 iterations where a randomly chosen item from each category is left-out. Performance is then averaged over iterations for each participant. For conditions where we train and test on orthogonal sets of data we simply train and test once on all of the data. In each case, we then perform one-sample t-test vs. chance performance (50%) to assess the statistical significance of the decoding accuracy. We apply this SVM to training and testing based on: (1) whether an unambiguous face vs. non-face was shown (decoding the stimulus itself); (2) whether an ambiguous face trial was subsequently remembered or forgotten, and (3) whether training on the perceptual distinction of unambiguous face vs. non-face transfers to testing on remembered vs. forgotten ambiguous faces, as expected if this memory difference is driven by whether the items are perceived as faces, and (4) whether training on the distinction of remembered vs. forgotten for (a) non-faces, and (b) unambiguous faces transfers to testing on remembered vs. forgotten ambiguous faces, as would be expected if there were general memory strength signals, like attention.

#### ERP Analysis

In addition to the classification analysis, we also conducted a planned analysis using event-related potentials (ERPs). In particular, we assessed the magnitude of the N170, a measure of face processing (Bentin, Allison, Puce, Perez, & McCarthy, 1996; Eimer, 2000; George, Evans, Fiori, Davidoff, & Renault, 1996), while participants encoded the images. Our main question of interest was whether this neural signature of category-specific processing would predict which ambiguous-face images were remembered and which were forgotten. In particular, since the amplitude of the N170 is indicative of which images participants’ see as a face (George, Jemel, Fiori, Chaby, & Renault, 2005; Hadjikhani, Kveraga, Naik, & Ahlfors, 2009; Liu et al., 2014), this component should provide a neural measure of category-specific processing for each image, which we hypothesized will predict which images will be memorable (Bower et al., 1975; Wiseman & Neisser, 1974). To assess this, ERPs were averaged separately for unambiguous face images, remembered-ambiguous face images, forgotten-ambiguous face images, and non-face images for each participant separately. ERPs were digitally low-pass filtered (−3dB cutoff at 25 Hz) and the mean amplitude of the N170 component was measured between 140 to 180 ms at two right posterior electrodes PO8 and P8 (Eimer, 2000) with respect to a 200-ms pre-stimulus period. The mean amplitudes were subjected to a repeated-measures analysis with image type as a within-subject factor. Planned pairwise comparisons were conducted to examine the N170 for unambiguous face images vs. non-face images, and remembered-ambiguous face images vs. forgotten-ambiguous face images.

## Results

### Behavioral results

We quantified memory performance primarily in terms of sensitivity (d’). As expected, participants remembered the unambiguous face images with the highest accuracy, followed by the ambiguous face images, and lastly the non-face images (see Figure 2A; *F*(2,46)=47, *p*<0.0001, |^2^=0.67). Pairwise comparisons revealed that all three conditions differed reliably from each other (unambiguous face vs. non-face: *t*(23)=8.19, *p*<0.0001, |^2^=0.75; unambiguous face vs. ambiguous face: *t*(23)=3.45, *p*=0.002, |^2^=0.34 ; ambiguous face vs. non-face: *t*(23)=6.28, *p*<0.0001, |^2^=0.63). These effects were accounted for more by a difference in hit rates (74.9%, 61.4% and 50.2%, respectively, for unambiguous-faces, ambiguous-faces and non-faces) than a difference in false alarm rates (24.3%, 18.4%, 24.5%, respectively).

**Figure 2.**
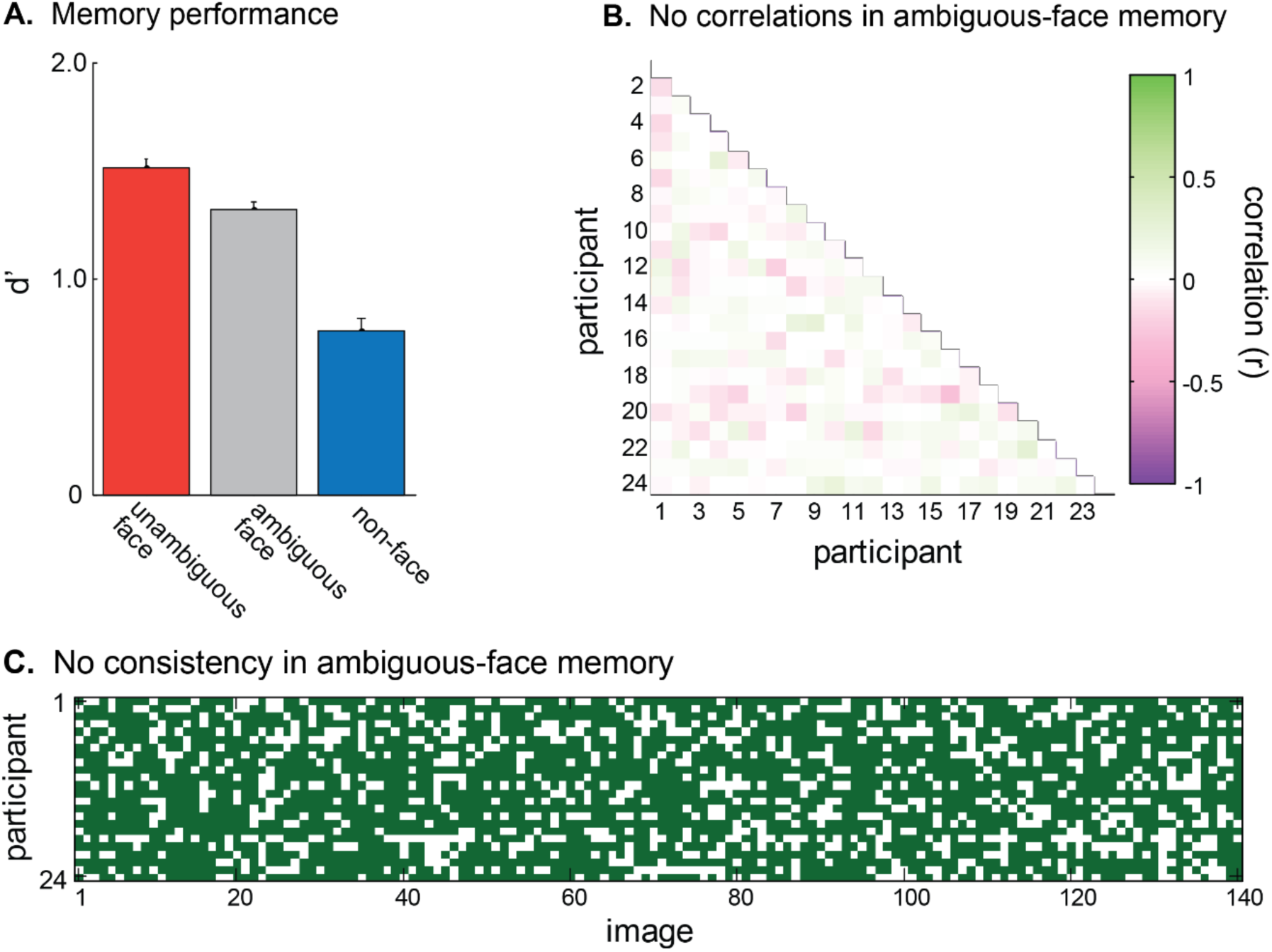
(A) Memory performance for unambiguous faces, ambiguous faces and non-faces, in terms of d’ (sensitivity). Participants were better able to remember unambiguous faces than non-faces, with ambiguous faces falling in between. Error bars represent ±1 S.E.M. (B) Correlations between participants in which ambiguous faces were correctly remembered. The lack of consistent correlation (mean r=0.02) indicates that different participants remembered different ambiguous-faces. (C) An alternative representation of this lack of consistency in which ambiguous-face images participants remembered, showing, for each of the 140 ambiguous-face images tested, which participants got the test trial correct (=green) or incorrect (=white). The lack of vertical columns indicates the lack of consistency.

We found that participants were extremely inconsistent in which ambiguous face images they found easiest to distinguish from new images (see Figure 2B, 2C). In particular, the correlation between participants in which images were correctly categorized as old/new at test is extremely low (*r*=0.02); less than 1% of the variance in which ambiguous faces were remembered was predicted by the images remembered by other participants (Figure 2B). This is visualized in Figure 2C by showing each of the ambiguous Mooney face images as a column and indicating in green when a participant correctly categorized it as old/new at test. The lack of consistency is shown by the lack of vertical column structure (e.g., participants do not consistently get the same images correct or incorrect). Thus, on average, different participants remembered different ambiguous-faces. This means that we can examine neural activity involved in successful memory encoding for ambiguous stimuli independent of the particular images.

### Decoding results

We first established a baseline for how well we could decode whether a stimulus currently being viewed is an unambiguous face or a non-face using single-trial EEG activity, effectively establishing an upper bound for decoding accuracy in our stimulus set. In particular, we trained an SVM to distinguish between unambiguous faces and non-faces based on EEG data averaged over 20ms bins at all electrodes during the encoding interval (100–600ms post-stimulus), and then tested this classifier on a leave-one-out hold-out set (see Methods). On average across participants, we found decoding accuracy of 61.8% (SEM: 1.8%), well above-chance (*t*(23)=6.63, *p*<0.001; see Figure 3A). Thus, despite the matched low-level features, we can decode whether a Mooney image that is currently being processed is a face or a non-face, suggesting that participants process face and non-face stimuli distinctly even when they are Mooney images (Kanwisher, Tong, & Nakayama, 1998; Hsieh, Vul & Kanwisher, 2010). Although this upper bound of decoding accuracy is rather low, this is expected given noisy single-trial EEG data and well-matched perceptual stimuli, in line with previous studies that have tried to decode subsequent memory or task performance in general using single-trial EEG activity (e.g., Höhne et al., 2016; Leydecker et al., 2014; Noh et al., 2014).

**Figure 3.**
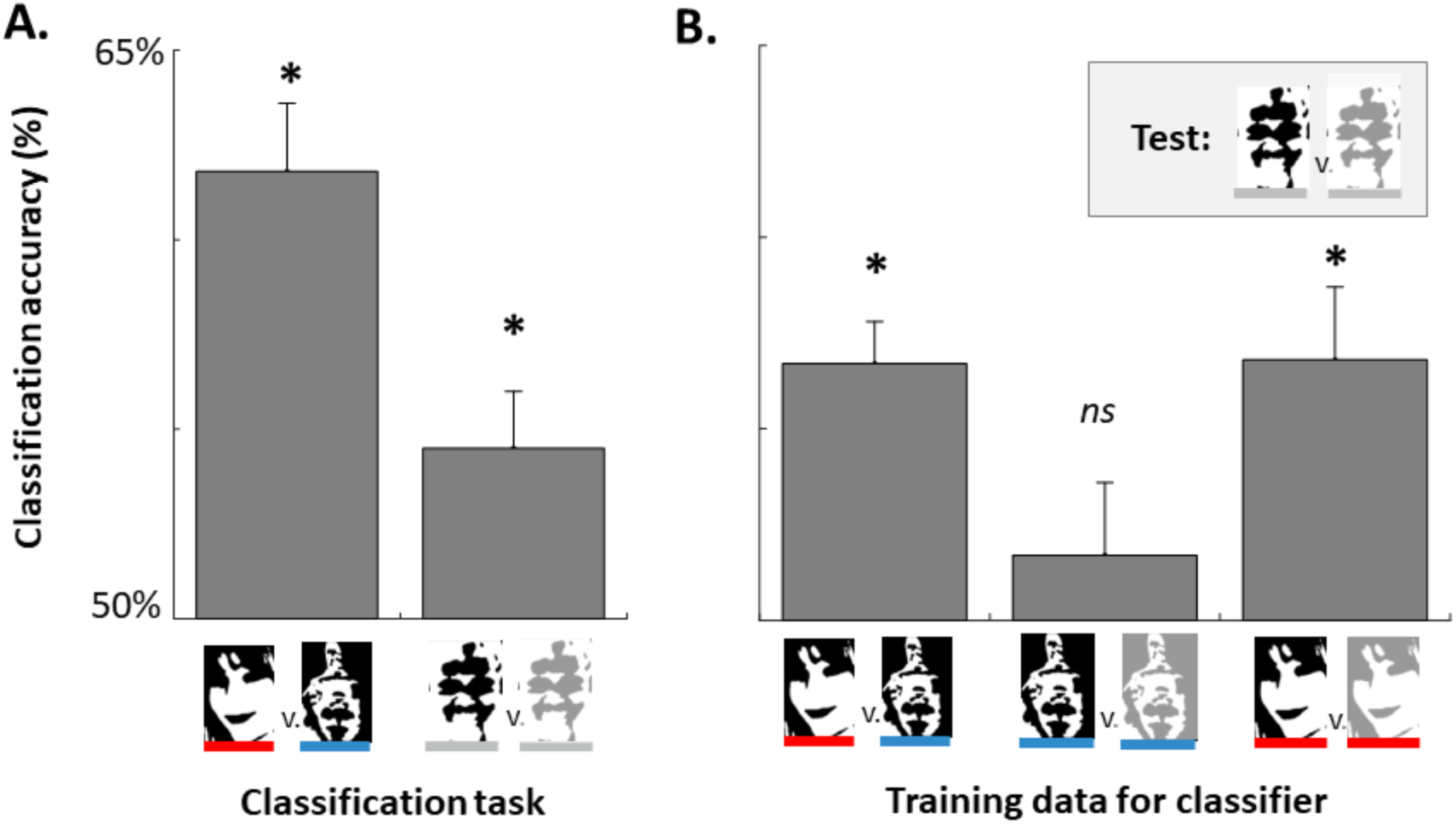
A) Decoding accuracy for a classifier applied to predicting at the time of encoding whether (Left) people are seeing a face vs. non-face and (Right) whether they will subsequently remember an ambiguous face stimulus (i.e., decoding based on encoding activity ambiguous-remembered faces vs. ambiguous-forgotten faces). B) Generalization tests for the classifier. To determine what predicts subsequent memory for ambiguous faces, we trained a classifier on other discriminations and asked whether it could predict subsequent memory for ambiguous faces. (Left) A classifier trained to discriminate between seeing a face vs. seeing a non-face predicted subsequent memory for ambiguous faces, such that images perceived as faces were more likely to be subsequently remembered. (Middle) A classifier trained to discriminate subsequently remembered vs. forgotten non-faces could not predict subsequent memory for ambiguous face stimuli. (Right) A classifier trained to discriminate subsequently remembered vs. forgotten unambiguous faces could predict subsequent memory for ambiguous face stimuli as well.

Our main question of interest was how participants encoded ambiguous face stimuli and whether ambiguous face stimuli that were subsequently remembered were processed distinctly from those that were subsequently forgotten. To assess this, we first examined whether subsequent memory could be decoded from the encoding related activity to ambiguous faces, and then, using a variety of generalization tests, asked what distinctions in the way the stimuli were processed might lead to this decoding.

We found that we could decode subsequent memory status for the ambiguous face stimuli. That is, we can decode from the signal at the time of encoding whether an ambiguous face will be subsequently remembered or subsequently forgotten (54.5%, SEM: 1.5%). Although the overall performance level is modest, it is highly consistent across observers (*t*(23)=2.91, *p*=0.008).

What drives this difference in subsequent memory for ambiguous faces? There are least two non-mutually exclusive possibilities: First, there is the possibility that the ability to decode subsequent memory is a result of the ambiguous face stimuli sometimes being recognized as a face vs. not (as shown in the behavioral results). Thus, the ambiguous face data may be made up of some encoding events that are similar to the unambiguous face trials (e.g., where participants saw a face) and others that are similar to the non-face trials (e.g., where participants did not see the face). If this were the case, then a classifier trained to distinguish between unambiguous faces and non-faces, without regard to memory performance should successfully generalize and without further training predict subsequent memory for ambiguous faces. That is, if the classifier believes the ambiguous face was "seen as a face", this should predict that the item was subsequently remembered. Indeed, using this approach results in a decoding accuracy of 56.7% (SEM: 1.1%) for distinguishing subsequently remembered vs. non-remembered ambiguous faces, significantly above-chance (*t*(23)=6.06, *p*<0.00001). In fact, this generalization performance was not significantly different than attempting to decode memory directly (as above, M=54.5% accuracy; *t*(23)=-1.30, *p*=0.207).

Another possibility that could help explain subsequent memory for ambiguous faces is a general subsequent memory effect. That is, independent of whether items are recognized as faces, some items may simply be better attended or further processed during encoding, resulting in a generic subsequent memory signal. To assess this possibility we trained a classifier to predict subsequent memory for non-face stimuli. We then asked if this classifier generalizes to predict subsequent memory for ambiguous faces. We did not find evidence consistent with a generic subsequent memory signal in decoding the single-trial EEG data (generalization performance: 51.7%, SEM: 1.9%, t(23)=0.90, p=0.38).

Lastly, we examined whether there was a more stimulus-specific subsequent memory signal, e.g., a signal of how strongly a stimulus was processed that was specific to face-like stimuli. To examine this, we next trained a classifier to predict subsequent memory for unambiguous face stimuli. We then asked if this classifier generalizes to predict subsequent memory for ambiguous faces. We found evidence consistent with this possibility (generalization performance: 56.8%, SEM 1.9%, t(23)=3.65, p=0.001).

Overall, our decoding results suggest that ambiguous faces that are processed *as faces* are subsequently remembered more often than those that elicit non-face-like activity at encoding. Thus, a classifier trying only to predict whether a stimulus is a face or non-face predicts subsequent memory for ambiguous faces even using only single-trial EEG data. In addition, we find that the same classifier that predicts the likelihood of an unambiguous face being remembered generalizes to predict the likelihood of an ambiguous face being remembered; but that the same is not true for a classifier trained to predict subsequent memory for non-faces. This is again consistent with the idea that subsequent memory for ambiguous faces is driven primarily by the strength of a face-specific response or a face-specific memory signal.

Importantly, this hypothesis is directly testable, because of the existence of a face-selective ERP component, the N170 (George, Jemel, Fiori, Chaby, & Renault, 2005; Hadjikhani, Kveraga, Naik, & Ahlfors, 2009; Liu et al., 2014). If the decoding of subsequent memory is largely driven by the strength of face-specific processing, we should be able to see the subsequent memory effect for ambiguous faces not only using single-trial decoding but also more selectively in the N170 component. This can help us localize the source of this decoding accuracy and pinpoint to what extent it is due to face-specific processing.

### ERP results

Since the amplitude of the N170 is indicative of which images participants’ see *as a face* (George, Jemel, Fiori, Chaby, & Renault, 2005; Hadjikhani, Kveraga, Naik, & Ahlfors, 2009; Liu et al., 2014), this component provides a useful neural measure of category-specific processing for each image, allowing us to directly test the extent to which category-specific processing drives subsequent memory even with stimuli that are perceptually identical on average.

We find that the mean amplitude of the N170 elicited during the encoding period differed reliably for unambiguous faces, remembered ambiguous faces, forgotten ambiguous faces and non-faces (F(3,69)=4.64, *p*=0.005, |^2^=0.17). As expected, the unambiguous face images elicited a larger N170 than the non-face images (t(23)=2.54, *p*=0.018, |^2^=0.22; see Figure 4A left), verifying that the N170 is sensitive to faces even in our Mooney images.

Our main question of interest was whether the N170 amplitude differed for subsequently remembered and subsequently forgotten ambiguous face images, even though across participants, different images were remembered or forgotten and thus the perceptual input was matched across the conditions. We found that the amplitude of the N170 did indeed predict memory performance for the ambiguous faces: For those images that participants later remembered correctly, the N170 was larger than the N170 for those images that participants later forgot (*t*(23)=2.63, *p*=0.015, |^2^=0.23; see Figure 4A right). Furthermore, the N170 elicited by ambiguous-remembered faces was not different than that elicited by unambiguous faces (p=0.95) but was larger than the N170 elicited by non-faces (*t*(23)=2.32, *p*=0.03, |^2^=0.19). Moreover, the N170 elicited by ambiguous-forgotten faces was not statistically different than that elicited by non-faces (*p*=0.67) but was smaller than the N170 elicited by unambiguous faces (*t*(23)=3.08, *p*=0.005, |^2^=0.29).

Thus, a very strong predictor of whether an image was later remembered by a particular participant was whether that image elicited category-specific brain activity during the encoding period, as indexed by the amplitude of the N170.

**Figure 4.**
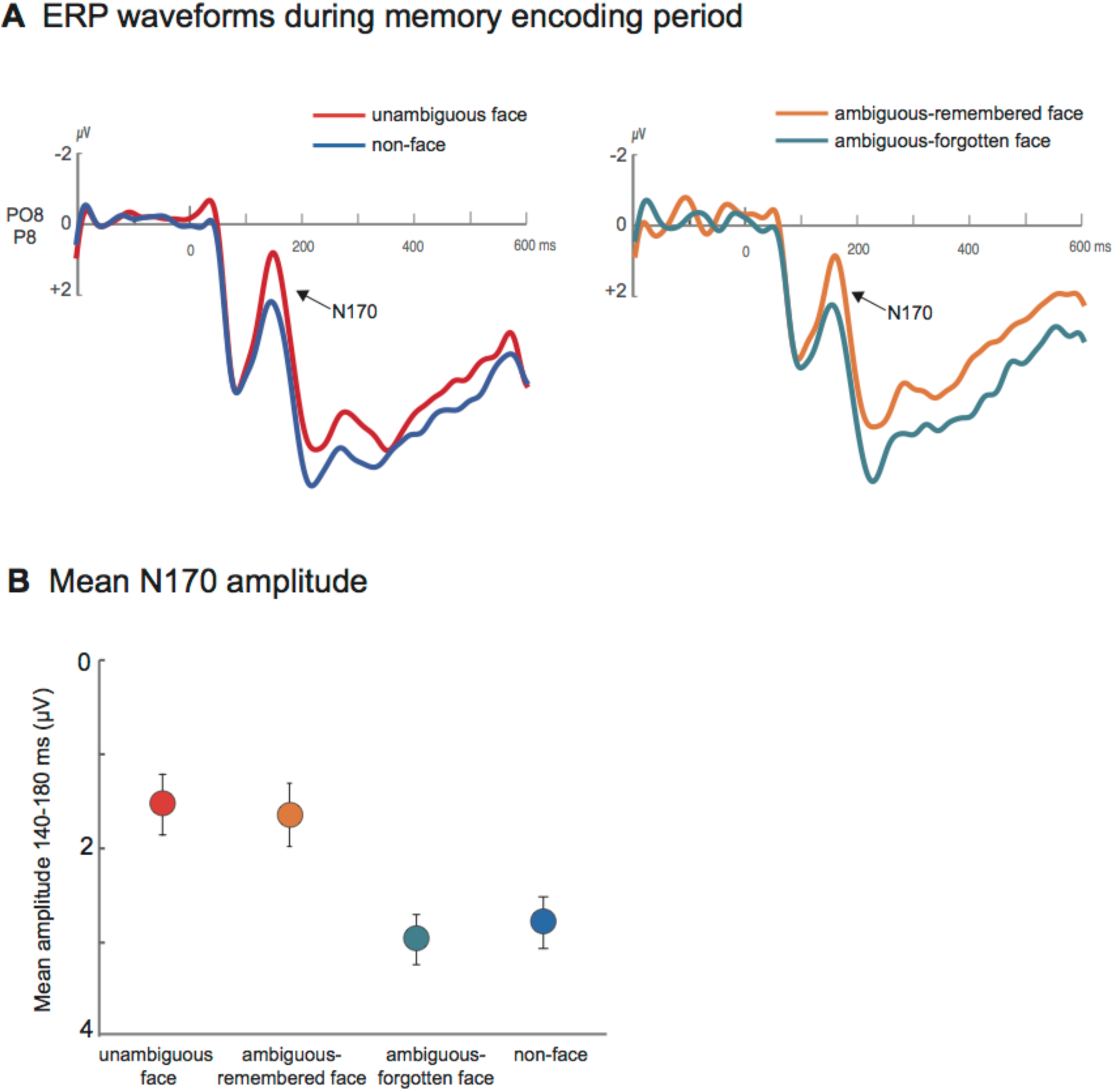
A) ERP waveforms during the memory encoding period. Left: the N170 is larger for unambiguous-faces relative to non-faces. Right: the N170 is also larger for ambiguous faces that were later remembered than for ambiguous faces that were later forgotten. B) Mean N170 amplitude for the 140–180ms time-window. Error bars represent ±1 S.E.M.

We find a large effect of subsequent memory on the size of the N170 for ambiguous-face images, which can be either seen as a face or not seen as a face by particular participants. We interpret the increased N170 for subsequently remembered items as evidence that items that are seen as a face are more likely to be remembered. To ensure that the amplitude difference measured during the N170 time interval does not simply reflect a general effect of subsequent memory performance that is unrelated to seeing the stimulus as a face, we also examined the mean amplitudes for remembered and forgotten unambiguous faces and non-faces.

For the unambiguous stimuli, the N170 was larger for faces than non-faces, regardless of subsequent memory performance (main effect of stimulus type: *F*(1,23)=5.83, *p*=0.02; no effect of memory accuracy: *p*=0.45; no interaction: *p*=0.80, see Figure 5), indicating that the N170 truly reflects whether participants see the images as a face and only predicts subsequent memory for the ambiguous-face stimuli.

**Figure 5.**
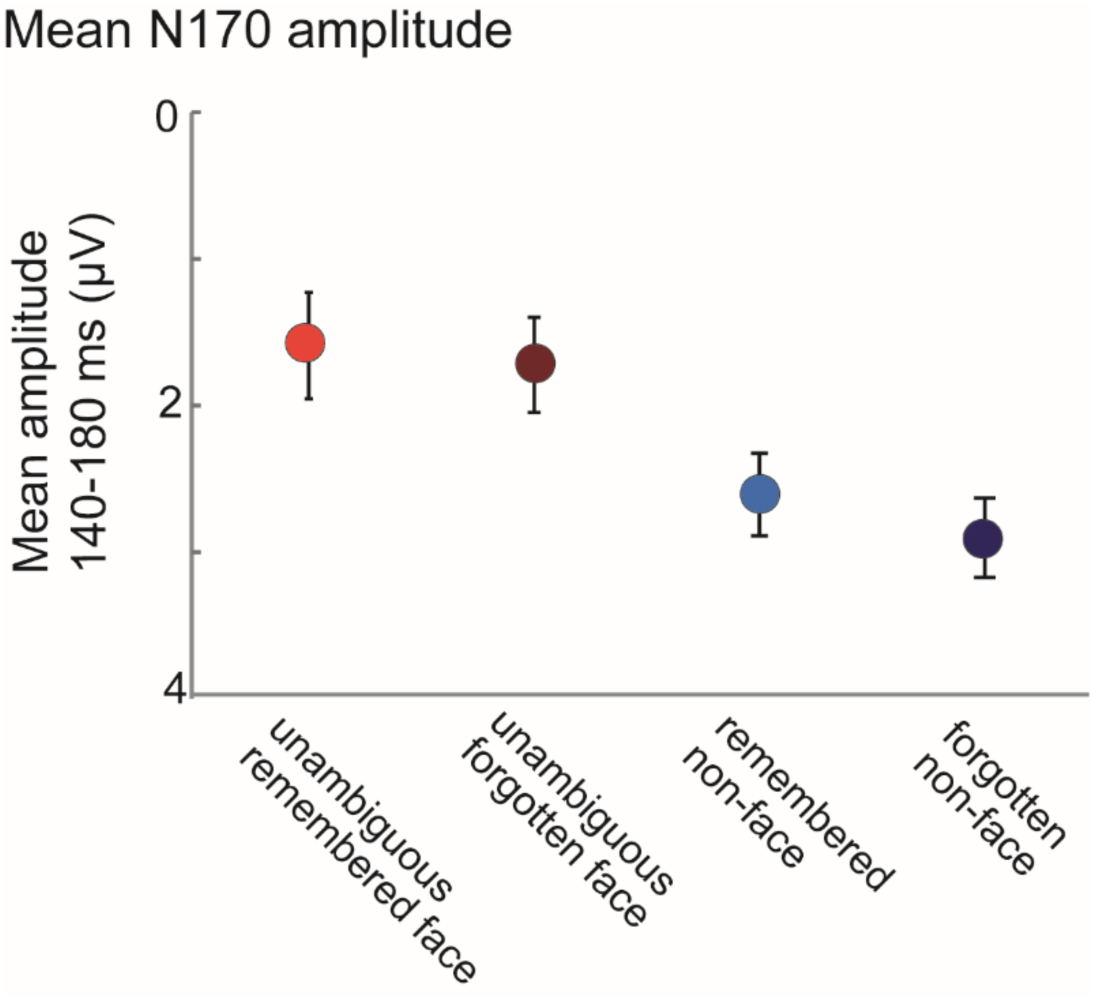
Regardless of whether they are subsequently remembered, unambiguous face stimuli always evoke a strong N170; whereas non-faces do not. This suggests that N170’s effect on subsequent memory is specific to the ambiguous face images, consistent with the role of meaning in visual memory. Error bars represent ±1 S.E.M.

## Discussion

We examined the role of meaning in visual memory. In particular, we asked participants to remember the visual details of an image – which exact set of black and white shapes they saw – and examined whether memory performance depended on whether this information was understood by the participants and connected to a meaningful interpretation (i.e. seeing it *as a face),* or whether the exact same perceptual input would be equally well remembered regardless of how well understood it was. We first showed behaviorally that participants are not consistent in which particular ambiguous face stimuli they remember. Thus, any neural prediction of subsequent memory performance for ambiguous face stimuli must result purely from the role of meaningfulness or other more general encoding factors like attention, not from the perceptual properties of the stimuli or differences in encoding strategies set up by the experimenter.

Using single-trial EEG activity during stimulus encoding, we were able to predict whether a particular ambiguous face stimulus was later remembered or forgotten. Critically, using a variety of generalization tasks, we found that the decoding accuracy cannot be explained by general differences at encoding that are shared between stimuli conditions, but rather is explained by whether a particular image elicited face-like activity at encoding for that particular person. This is further confirmed by the additional analysis of the face-selective N170 component in the ERP waveform which is unambiguously related to whether a stimulus was seen as a face or not.

Previous studies have shown that electrophysiological brain signals during stimulus encoding differ on average for items that are later remembered vs. forgotten (Sanquist et al., 1980; Paller & Wagner, 2002). These differences are often apparent in amplitude or latency modulations of the ERP (Paller, Kutas, & Mayes, 1987; Paller, McCarthy, & Wood, 1988; Fukuda & Woodman, 2015) as well as distinct oscillations ranging from changes in the alpha, theta or gamma rhythms (Hanslmayr, Spitzer, & Bauml, 2009; Klimesch et al., 1996; Osipova et al., 2006; Staudigl & Hanslmayr, 2013). A few recent studies have also shown that single-trial EEG activity can predict subsequent memory by combining pre-stimulus and stimulus-locked activity during encoding, reaching similar decoding accuracies as in the present study (59.6%; Noh et al., 2013; see also Tan, Smitha, & Vinod, 2015; Sun, Qian, Chen, Wu, Luo, & Pan, 2016). However, in all of these studies, differences in encoding-related brain activity could be due to differences in the perceptual input (the same images tend to be remembered by the same people), differences in familiarity of the stimuli, distinct encoding strategies, or the attentional state of the observer at encoding (Klimesch, 2012; Dubé et al., 2013; Hanslmayr & Staudigl; 2014). In contrast, our results uniquely indicate that differential processing of the exact same stimuli, and in particular the extent to which these stimuli are processed in a category-specific and meaningful manner, predicts subsequent memory independent of other attentional or perceptual factors.

Our results fit with existing data showing that category-specific brain regions are involved in encoding information into long-term memory (Kuhl, Rissman, & Wagner, 2012; Prince, Dennis, & Cabeza, 2009), but go beyond this data by carefully controlling for the perceptual properties of remembered and forgotten stimuli and by contrasting meaningful vs. non-meaningful stimuli, as opposed to examining only memory for meaningful information. In particular, in previous data showing category-specific brain responses predict subsequent face memory, the perceptual properties of the stimuli are not matched across subsequent memory effects, and there are likely systematic differences between the faces that are remembered and those that are not. For example, same-race faces are both better remembered and provoke more FFA activity during perception than other-race faces (Golby, Gabrieli, Chiao, & Eberhardt, 2001); and particular characteristics of faces predict which ones are better remembered in a way that is reliable across individuals (Bainbridge, Isola, & Oliva, 2013; Khosla, Bainbridge, Torralba, & Oliva, 2013). By contrast, our data show that perceiving an image as meaningful – as reflected in the category-specific brain response - rather than treating it as an unrecognizable set of shapes, is directly related to better subsequent memory, even for identical images. This result demonstrates the important role of meaning in visual memory.

Our data broadly suggest an important role for the meaning of a stimulus, rather than its perceptual properties, in explaining detailed visual memory performance (Bower et al., 1975; Konkle, Brady, Alvarez, & Oliva, 2010; Koutstaal et al., 2003; Wiseman & Neisser, 1974). Why does meaning play a crucial role in visual memory? It has long been argued that memory is semantically organized (e.g., in spreading activation models; Collins & Loftus, 1975), and that both elaborative encoding and building strong retrieval cues is critical to successful long-term memory. In particular, it is now thought that many items that are lost in long-term memory are likely not forgotten entirely; instead, retrieval fails because no distinctive retrieval cue allows access to the item (Wixted, 2004). A meaningful interpretation of an image may allow for a more elaborative initial encoding, creating more or stronger paths to access the memory (Bradshaw & Anderson, 1982). Thus, meaning may act as a “hook” to allow the retrieval of even visual details by creating specific retrieval cues (Konkle et al., 2010).

In terms of neural representation, meaning may play a role in so far as more neural populations can be recruited for meaningful stimuli than non-meaningful stimuli, resulting in more robust memory representations that are less limited by interference (e.g., Cohen et al. 2014). Because more relevant neural populations are available to support memory traces for faces and other meaningful stimuli than for non-sense blobs, these stimuli may have either more robust memory traces, or these traces may be more distinctive from each other and thus less likely to result in interference in memory. This is broadly consistent with the role of interference in long-term memory (e.g., Wixted, 2004). For example, cognitive studies have revealed that even with matched perceptual variation, categories that have a wider variety of conceptual information result in reduced interference between items in that category (Konkle et al. 2010), compatible with the idea that items from more conceptually broad categories may have more distinct memory traces.

Our data provide a neural analog and an important refinement of a classic result by Wiseman and Neisser (1974), who used a similar paradigm to the one we used in the current study. Wiseman and Neisser showed that when participants are shown faces for extended durations (5 seconds) and are explicitly asked to indicate whether an image is a face or not, they later remember the face images more accurately. In their study, which images were recognized as a face by different participants was not random, leaving open the possibility that the low-level visual features of the images were driving memory performance rather than solely whether they connected to meaningful interpretations for the participants. In addition, their long presentation times – required to perform the explicit face categorization task – means that subsequent memory effects could be caused by the fact that participants enjoy looking at faces more than unrecognizable shapes and so spend a larger portion of the 5s encoding window focusing on stimuli they recognize as faces. In the current study we were able to pick up the brain response spontaneously elicited by briefly presented ambiguous-face images and to control for perceptual features in the remembered and non-remembered stimuli. Our results thus provide a much stronger test of the role of meaningfulness in subsequent memory.

Overall, the current data provide strong evidence for the involvement of meaning in visual memory, even in the absence of any perceptual differences between stimuli that evoke meaning and those that do not. Stimuli that evoke category-specific brain activity during encoding are better remembered than identical stimuli that do not evoke this brain activity. This provides a strong dissociation between perceptual factors and meaning, and suggests that even detailed visual memory is strongly supported by the meaning of a stimulus.

## Author Contributions

T.F.B. and V.S.S. designed and conducted the experiment and analyzed the data. T.F.B., G.A.A. and V.S.S. contributed to the writing of the manuscript and contributed to the motivation and overall research question.

## References

Bainbridge, W., Isola, P., & Oliva, A. (2013). The intrinsic memorability of face photographs. Journal of Experimental Psychology: General, 142(4), 1323.

Bartlett, F. C. (1932). Remembering: A study in experimental and social psychology. New York: Macmillan.

Bentin, S., Allison, T., Puce, A., Perez, E., & McCarthy, G. (1996). Electrophysiological Studies of Face Perception in Humans. Journal of Cognitive Neuroscience.

Bower, G. H., Karlin, M. B., & Dueck, A. (1975). Comprehension and memory for pictures. Memory & cognition, 3(2), 216–220.

Bradshaw, G. L., & Anderson, J. R. (1982). Elaborative encoding as an explanation of levels of processing. Journal of Verbal Learning and Verbal Behavior, 21(2), 165–174.

Cohen, G. (1990). Why is it difficult to put names to faces? British Journal of Psychology, 81, 287–297.

Collins, A., & Loftus, E. (1975). A spreading-activation theory of semantic processing. Psychological review, 82(6), 407.

Cox, D. D., & Savoy, R. L. (2003). Functional magnetic resonance imaging (fMRI)“brain reading”: detecting and classifying distributed patterns of fMRI activity in human visual cortex. Neuroimage, 19(2), 261–270.

Daselaar, S. M., Prince, S. E., & Cabeza, R. (2004). When less means more: deactivations during encoding that predict subsequent memory. Neuroimage, 23(3), 921–927.

Delorme, A., & Makeig, S. (2004). EEGLAB: An open source toolbox for analysis of single-trial EEG dynamics including independent component analysis. Journal of Neuroscience Methods, 134(1), 9–21.

Eimer, M. (2000). The face-specific N170 component reflects late stages in the structural encoding of faces. Neuroreport, 11(10), 2319–2324.

Fukuda, K., … Woodman, G. F. (2015). Predicting and improving recognition memory using multiple electrophysiological signals in real time. Psychological science, 26(7), 1026–1037.

George, N., Evans, J., Fiori, N., Davidoff, J., & Renault, B. (1996). Brain events related to normal and moderately scrambled faces. Cognitive Brain Research, 4(2), 65–76.

George, N., Jemel, B., Fiori, N., Chaby, L., & Renault, B. (2005). Electrophysiological correlates of facial decision: Insights from upright and upside-down Mooney-face perception. Cognitive Brain Research, 24(3), 663–673.

Golby, A. J., Gabrieli, J. D., Chiao, J. Y., & Eberhardt, J. L. (2001). Differential responses in the fusiform region to same-race and other-race faces. Nature neuroscience, 4(8), 845–850.

Hadjikhani, N., Kveraga, K., Naik, P., & Ahlfors, S. P. (2009). Early (M170) activation of face-specific cortex by face-like objects. Neuroreport, 20(4), 403–407.

Hanslmayr, S., Spitzer, B., & Bäuml, K. H. (2008). Brain oscillations dissociate between semantic and nonsemantic encoding of episodic memories. Cerebral cortex, 19(7), 1631–1640.

Hanslmayr, S., & Staudigl, T. (2014). How brain oscillations form memories—a processing based perspective on oscillatory subsequent memory effects. Neuroimage, 85, 648–655.

Hollingworth, A. (2004). Constructing visual representations of natural scenes: The roles of short- and long-term visual memory. Journal of Experimental Psychology: Human Perception and Performance, 30, 519–537.

Höhne, M., Jahanbekam, A., Bauckhage, C., Axmacher, N., & Fell, J. (2016). Prediction of successful memory encoding based on single-trial rhinal and hippocampal phase information. Neuroimage, 139, 127–135.

Hsieh, P. J., Vul, E., & Kanwisher, N. (2010). Recognition alters the spatial pattern of FMRI activation in early retinotopic cortex. Journal of neurophysiology, 103(3), 1501–1507.

Isola, P., Xiao, J., Torralba, A., & Oliva, A. (2011, June). What makes an image memorable?. In Computer Vision and Pattern Recognition (CVPR), 2011 IEEE Conference on (pp. 145–152). IEEE.

Kanwisher, N., Tong, F., & Nakayama, K. (1998). The effect of face inversion on the human fusiform face area. Cognition, 68(1), B1–B11.

Khosla, A., Bainbridge, W. A., Torralba, A., & Oliva, A. (2013). Modifying the memorability of face photographs. Proceedings of the IEEE International Conference on Computer Vision (pp. 3200–3207).

Konkle, T., Brady, T. F., Alvarez, G. A., & Oliva, A. (2010). Conceptual distinctiveness supports detailed visual long-term memory for real-world objects. Journal of Experimental Psychology: General, 139(3), 558. American Psychological Association.

Koutstaal, W., Reddy, C., Jackson, E. M., Prince, S., Cendan, D. L., & Schacter, D. L. (2003). False recognition of abstract versus common objects in older and younger adults: Testing the semantic categorization account. Journal of experimental psychology. Learning, memory, and cognition, 29(4), 499–510.

Kuhl, B. A., Rissman, J., & Wagner, A. D. (2012). Multi-voxel patterns of visual category representation during episodic encoding are predictive of subsequent memory. Neuropsychologia, 50(4), 458–469.

Leydecker, A., Biebmann, F., & Fazli, S. (2014, June). Single-trials ERPs predict correct answers to intelligence test questions. In Pattern Recognition in Neuroimaging, 2014 International Workshop on (pp. 1–4). IEEE.

Liu, J., Li, J., Feng, L., Li, L., Tian, J., & Lee, K. (2014). Seeing Jesus in toast: Neural and behavioral correlates of face pareidolia. Cortex, 53(1), 60–77.

Lopez-Calderon, J., & Luck, S. J. (2014). ERPLAB: an open-source toolbox for the analysis of event-related potentials. Frontiers in human neuroscience, 8(April), 213.

McWeeny, K., Young, A., Hay, D., & Ellis, A. (1987). Putting names to faces. British Journal of Psychology, 78, 143–146.

Mooney, C. M. (1957). Age in the development of closure ability in children. Canadian journal of psychology, 11(4), 219–226.

Noh E, Herzmann G, Curran T, de Sa VR. Using single-trial EEG to predict and analyze subsequent memory. NeuroImage. 2014; 84:712–723.

Paller, K. A., Kutas, M., & Mayes, A. R. (1987). Neural correlates of encoding in an incidental learning paradigm. Electroencephalography and clinical neurophysiology, 67(4), 360–371.

Paller, K. A., McCarthy, G., & Wood, C. C. (1988). ERPs predictive of subsequent recall and recognition performance. Biological psychology, 26(1–3), 269–276.

Prince, S. E., Dennis, N. A., & Cabeza, R. (2009). Encoding and retrieving faces and places: Distinguishing process- and stimulus-specific differences in brain activity. Neuropsychologia, 47(11), 2282–2289.

Sanquist TF, Rohrbaugh JW, Syndulko K, Lindsley DB. Electrocortical signs of levels of processing: perceptual analysis and recognition memory. Psychophysiology. 1980; 17(6):568–576.

Schwiedrzik CM, Melloni L, Schurger A (2018) Mooney face stimuli for visual perception research. PLoS ONE, 13(7): e0200106.

Sun, X., Qian, C., Chen, Z., Wu, Z., Luo, B., & Pan, G. (2016). Remembered or Forgotten?—An EEG-Based Computational Prediction Approach. PloS one, 11(12), e0167497.

Tan, Z. H. E., Smitha, K. G., & Vinod, A. P. (2015, October). Detection of Familiar and Unfamiliar Images Using EEG-Based Brain-Computer Interface. In Systems, Man, and Cybernetics (SMC), 2015 IEEE International Conference on(pp. 3152–3157). IEEE.

Wagner, A. D., Schacter, D. L., Rotte, M., Koutstaal, W., Maril, A., Dale, A. M., … & Buckner, R. L. (1998). Building memories: remembering and forgetting of verbal experiences as predicted by brain activity. Science, 281(5380), 1188–1191.

Wiseman, S., & Neisser, U. (1974). Perceptual organization as a determinant of visual recognition memory. The American Journal of Psychology, 87(4), 675–681. JSTOR.

Wixted, J. T. (2004). The psychology and neuroscience of forgetting. Annual review of psychology, 55(1), 235.

